# Molecular phylogeny and diversification timing of the Nemouridae family (Insecta, Plecoptera) in the Japanese Archipelago

**DOI:** 10.1101/440529

**Authors:** Maribet Gamboa, David Muranyi, Shota Kanmori, Kozo Watanabe

**Affiliations:** Department of Civil and Environmental Engineering, Ehime University, 790-0871 Matsuyama, Japan; Deparment of Zoology, Plant Protection Institute Centre for Agricultural Research, Hungarian Academy of Sciences, Herman Ottó u. 15, H-1022 Budapest, Hungary

**Keywords:** Stoneflies, Japan, divergence time, molecular phylogeny, speciation

## Abstract

The generation of the high species diversity of insects in Japan was profoundly influenced by the formation of the Japanese Archipelago. We explored the species diversification and biogeographical history of the Nemouridae family in the Japanese Archipelago using mitochondrial DNA and nuclear DNA markers. We collected 49 species among four genera: *Indonemoura*, *Protonemura*, *Amphinemura* and *Nemoura* in Japan, China, South Korea and North America. We estimated their divergence times—based on three molecular clock node calibrations—using Bayesian phylogeography approaches. Our results suggested that Japanese Archipelago formation events resulted in diversification events in the middle of the Cretaceous (<120 Ma), speciation in the Paleogene (<50 Ma) and intra-species diversification segregated into eastern and western Japan of the Fossa Magna region at late Neogene (20 Ma). The *Indonemoura* samples were genetically separated into two clades—that of Mainland China and that of Japan. The Japanese clade clustered with the Nemouridae species from North America, suggesting the possibility of a colonisation event prior to the formation of the Japanese Archipelago. We believe that our results enhanced the understanding both of the origin of the species and of local species distribution in the Japanese Archipelago.

## Introduction

The East Asian region—and in particular, the Japanese Archipelago—is considered to have high insect biodiversity [1], [2]. The high degree of Japanese insect biodiversity is a result of several mechanisms—in particular, the complex geological history. The Japanese Archipelago originated in the middle of the Miocene [3] as an independent formation of eastern and western Japanese landmasses. Extensive geographical changes and large-scale climatic changes throughout the islands facilitated the subsequent connection and disconnection of Japanese landmasses from the Eurasian continent, and the formation of tectonic lines (as the median tectonic line, MTL; and the Itoigawa-Shizuoka tectonic line, ISLT) [3], [4], [5]. These geological events—allowing for the colonisation of insects from the continent and their subsequent diversification as endemic lineages (i.e. new species)—contributed substantially to the high diversity of insects in Japan [2].

The process of species diversification has been intensively explored through phylogeographical approaches [6], [7]. These approaches have allowed for the observation of the historical process responsible for the current geographical distribution of individuals [6]. Molecular approaches to phylogeographic studies, using specific genes—such as mitochondrial DNA (mtDNA) or nuclear DNA (nDNA)—allow for a better understanding of species diversity by resolving complex taxonomic groups of species (for instance, cryptic species and species groups) [7]. Molecular phylogeography has provided valuable insights into the historical process of Japanese Archipelago formation underlying insect diversification. Previous studies identified genetic differentiation within species between the Japanese landmasses and the Eurasian continent (for instance, the mayflies *Isonychia japonica* [8]; caddisflies *Palaeagapetus* spp. [9]; and beetles *Ohomopterus* spp. [10] and the Carabina subtribe [11]). Dispersal events via land bridges (islands between continents) from the Eurasian continent to the Japanese Archipelago (of, for instance, the orthopteran *Locusta migratoria*, [12]; mayflies *Ephron* spp., [13]) or, in reverse, from the Japanese Archipelago to the Eurasian continent (of, for instance, water bugs *Appasus* spp., [14]) were additionally identified before, during and after the formation of the Japan Archipelago.

Aquatic insects have advantages in the studies of phylogeography, as their specialised ecological requirements and habitat range make aquatic insect species susceptible to geological changes. Among the Plecoptera order [15], the family Nemouridae is one of the largest and most dominant aquatic insect groups. The family comprises 20 genera and more than 400 species distributed throughout the Northern Hemisphere and across the equator in the Sunda Archipelago [16]. Several genera of the Nemouridae family have distinct disjunctions in their distribution [15]. For example, *Ostrocerca, Prostoia* and *Soyedina* were found in both the extreme western and the extreme eastern regions of North America, but they were absent in the central area [17], [18]. Similar disjunctive distributions were also observed among *Protonemura*, *Indonemoura*, *Sphaeronemoura* and *Illiesonemoura* in the Palaearctic region [19], the western and eastern Himalayan ranges [20] and North and South India [15]. *Podmosta* and *Zapada* are two interesting cases distributed across the Nearctic region and East Asia [21], [22]. Previous studies have suggested that their current habitat distribution could be associated with mountain formation and land bridges. In Japan, the Nemouridae family is widely distributed with four genera [23] —*Indonemoura*; *Protonemura*; *Amphinemura*; and *Nemoura.* To date, 30 *Nemoura* species, 17 *Amphinemura* species, 12 *Protonemura* species and 1 *Indonemoura* species have been reported in Japan [16]. However, their evolutionary history in the Japanese Archipelago remains unknown.

We studied the molecular phylogeny of the aquatic insect Nemouridae (Plecoptera) in the Japanese Archipelago with comprehensive genera-level sampling using mitochondrial cytochrome c oxidase 1 (*cox1*) and nuclear histone 3 (*H3*) markers. We hypothesised that the Nemouridae family diversification could be linked to the geological formation of the Japanese Archipelago. Therefore, we estimated the phylogenetic relationships among Nemouridae species and genera with reference to their historical biogeography. We focused on geographic events of Japanese Archipelago formation and their influence on the divergence time among the genera and species using a combination of fossil records and the Archipelago formation history. Furthermore, to estimate the historical process of the phylogeography of Nemouridae in Japan, we compared the phylogenetic relationships among the specimens from South Korea, China and North America, that are assumed to be the potential sources of Japanese Nemouridae because of the geological formation history of the Japanese Archipelago.

## Material and methods

### Study sites and sample collection

Our sampling sites in Japan comprised 32 sampling sites on Hokkaido Island, 83 on Honshu Island and 27 on Shikoku Island. None of the Nemouridae species was found on Hokkaido Island during sampling. All species reported from Hokkaido are known to occur on either Honshu or Shikoku Islands. Herein, we only reported on the sampling sites wherein specimens where found. We collected 20, 7, 8 and 1 species of the genera *Nemoura*, *Amphinemura*, *Protonemura* and *Indonemoura*, respectively, on 110 sampling sites in Japan (Fig 1, S1 Table). Additionally, 14 species distributed from 8 sampling sites of Mainland China and 2 of South Korea (S1 Table, Fig 2) and 100 specimens of the three species *Zapada columbiana*, *Z. cinctipes* and *Podmosta delicatula* (subfamily Nemourinae) collected from 15 sampling sites of North America (western United States of America and Alaska) were included in our analysis. We added these samples from outside of Japan because of their geographical proximity to the Japanese Archipelago and their geological formation histories.

**Fig 1.**
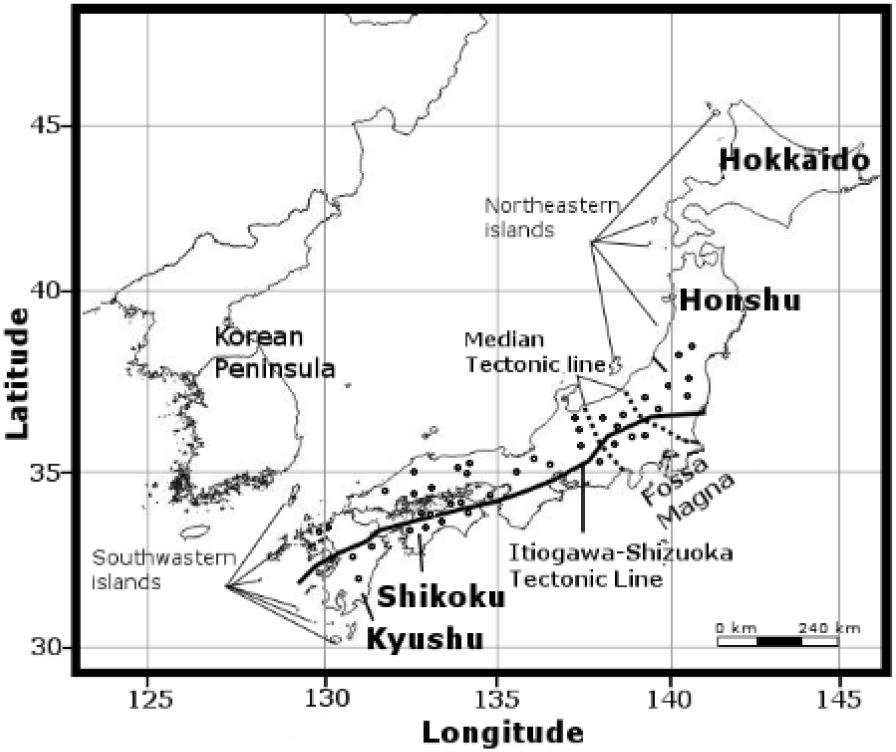
The Japanese islands and distribution of 110 sampling sites from where Nemouridae samples were collected (open circles).

**Fig 2.**
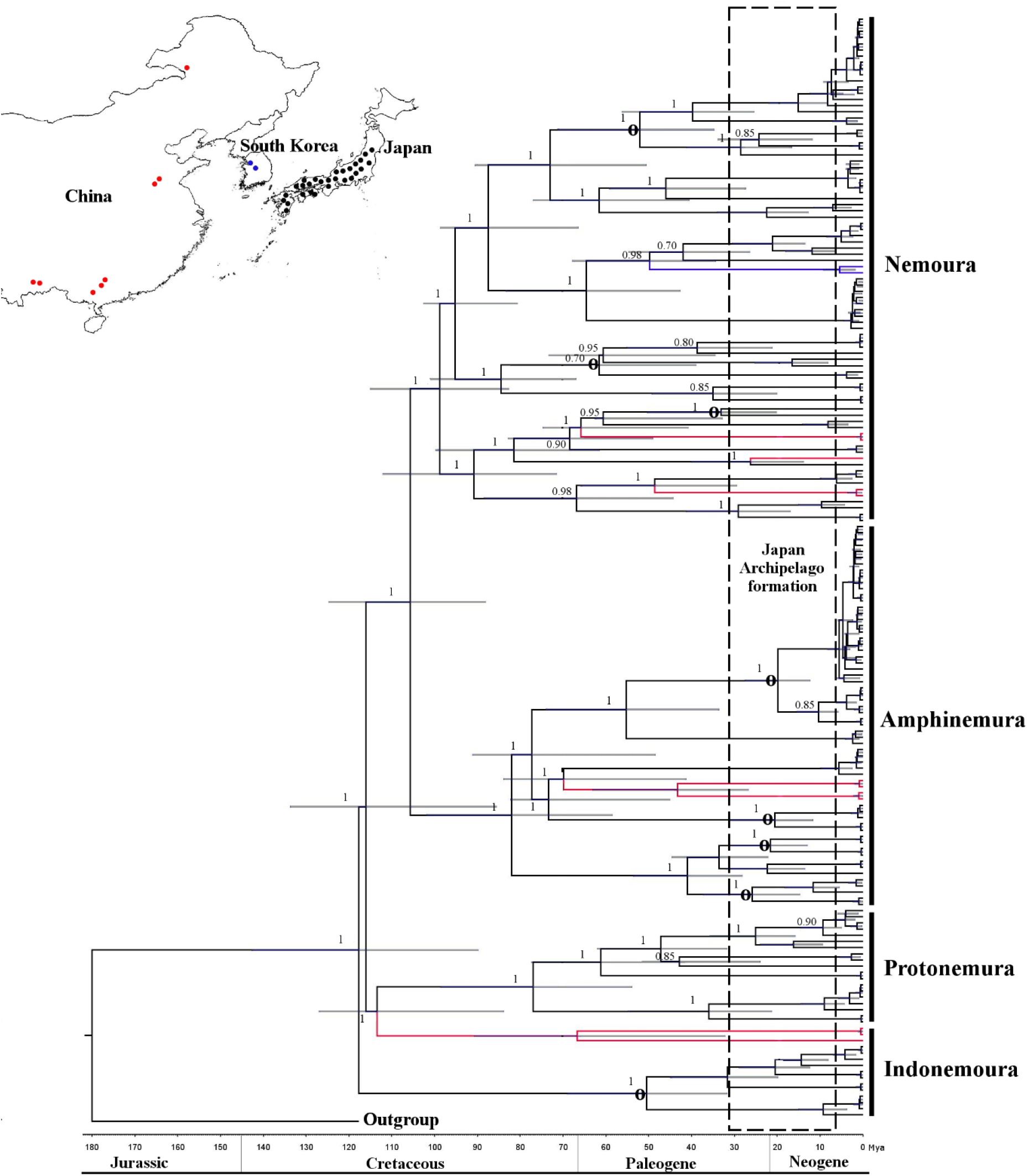
Concatenated Bayesian phylogeny (*cox1* + *H3*) of the East Asian Nemouridae family. The phylogenetic tree nodes were calibrated using 180 Ma based on fossil records + 15 to 30 Ma based on the Japanese Archipelago formation. Calibration and geological time are shown at the bottom of the tree. A 95% HPD is indicated as a horizontal grey bar and posterior probabilities are shown for each node. Circle symbol (**○**) in the nodes indicates intra-species diversification based on the eastern and western Japanese boundaries of the Fossa Magna region. Inserted upper map shows sample site locations for Japan (black), China (red) and South Korea (blue) as dots. Colour branches indicate sample location distribution as shown in the map.

We collected adult insects using hand nets around riversides. We stored samples in 80% ethanol in the field, and replaced the ethanol with fresh 99.5% ethanol after morphological identification. We identified individuals according to the taxonomical keys of [21], [23], [24], [25], [26], [27], [28], [29], [30], [31] and [32]. Undescribed species resulted in our studied were based on our taxonomic expertise and inconclusive taxonomic keys.

### DNA extraction, amplification and sequencing

We genetically analysed a total of 289 individuals, out of which 189 were from East Asia (males, 97; females, 92) and 100 were from North America (males, 92; females, 8). We extracted genomic DNA individually using DNeasy tissue kits (Qiagen GmbH, Hilden, Germany), following the manufacturer’s instructions. We amplified a 658-bp fragment of mtDNA *cox1* using LCO-1490 and HCO-2198 primers [33] with an annealing temperature of 38°C and 40 PCR cycles. Further, we amplified a 328-bp fragment of nDNA marker histone 3 (*H3*) using the universal primers H3F and H3R [34] with an annealing temperature of 58°C and 40 PCR cycles. We purified the PCR products using the QIAquick PCR Purification Kit (Qiagen GmbH, Hilden, Germany) and sequenced them in both directions using the same primers as mentioned above. *Cox1* and *H3* sequences were sequenced by Eurofins Operon (Tokyo, Japan). All sequence data reported here have been deposited in GenBank.

### Sequence analysis

We assembled and edited forward and reverse sequences using CodonCode Aligner v 3.5 (Codon Code Corporation, Dedham, USA). All sequences were aligned using ClustalW (https://www.genome.jp/tools-bin/clustalw) [35]. We calculated the genetic diversity by the number of polymorphic sites, number of haplotypes and both mean nucleotide substitution rate (i.e individuals within species) and pairwise nucleotide substitution rate (i.e between species), with the Kimura 2- parameter model. We performed all analyses using DnaSp v5.10 [36]. All analyses were performed for *cox1* and h3 separately.

All sequences of the mtDNA and nDNA markers were compared with the NCBI nucleotide database using blastn queries (http://blast.ncbi.nlm.nih.gov) to corroborate species identification (DNA barcoding, similarity > 98%) and to discard possible sequence errors.

### DNA species delimitation

To corroborate the morphological species identification match with our molecular data, we implemented a DNA species delimitation analysis. Putative DNA species were delineated using the General Mixed Yule Coalescent model (GMYC; [37]). An ultrametric gene tree of *cox1* gene was constructed using BEST v1.8.3 [38], and the GMYC analysis was performed using the splits package [39] in R ver. 3.3 (R Core Team).

### Molecular clock analysis

We estimated the evolutionary history of the family in the Japanese Archipelago according to the timing of the divergence of the lineages. For this estimation, we implemented a Bayesian phylogenetic analysis in combination with a molecular clock analysis using BEAST v.2.4.4 [40] with *Zwicknia bifrons* (Capniidae) as a outgroup for *cox1* and *H3* (own sequences) separately. This outgroup was selected owing to their close phylogenetic relationship with Nemouridae [41], [42]. To observed the divergence time, we adopted a relax clock model [43] following a log normal distribution, and calibrated the phylogenetic tree nodes using three types of molecular clock analysis. The first calibration was based on fossil records of the Nemouridae family [44]. We calibrated the nodes at 180 million years ago (Ma) and adjusted the parameters with a standard deviation of 20 Ma, as suggested in previous study [45] for a 95% highest posterior density (HPD). For this analysis, we implemented a fossilised birth death model [46] for tree prior parameter. The second calibration was based on the Japanese Archipelago formation events dated from 15 to 30 Ma [3]. We applied several calibrations from 15, 20, 25 and 30 Ma at all nodes representing taxonomic species. All calibrations were adjusted to 5 Ma as a standard deviation for a 95% HPD and a fossilised birth death model [46] for tree prior parameter. Lastly, the third calibration was the time to the most recent common ancestor (TMRCA) to observe species diversification patterns based on the mean substitution rate of *cox1*. Using a Yule model tree prior parameter [47], we applied the substitution rate for insect *cox1* of 1.5% [48] and 3.54% [49] per million years for a 95% HPD.

For all branch age calibrations (namely, fossil, biogeographic and mtDNA substitution rate), we performed MCMC for 50 million generations, and log dating trees (BEAST parameters) for every 5000 generations. We tested the output files for convergence after removing a 10% burn-in by examining the effective sampling size using Tracer v1.5 [50]. We pooled the four resulting output trees from biogeographical calibration analysis into a single tree. We then pooled the resulting single tree from biogeographical branch calibration and the single tree from fossil calibration analyses into a single tree. We performed all pooling analyses using Log Combiner v1.6.1 (BEAST package) summarised with Tree Annotator (BEAST package) and visualised using FigTree v1.3.1 [51]. We performed the analyses for *cox1* and *H3* separately. The incongruence length difference test (ILD) [52] was conducted to test the congruence of tree topologies between *cox1* and *H3* using Tree Analysis Technology (TNT) [53]. ILD test revealed no significant differences in terms of the Bayesian tree topologies between *cox1* and *H3* (P = 0.8); therefore, both markers were polled into a single tree for further analysis.

### Phylogenetic analysis between Nemoura from Japan and North America

To observe the phylogenetic relationship between Nemouridae from Japan and North America (*Zapada columbiana*, *Z. cinctipes* and *Podmosta delicatula*), we analysed the maximum likelihood (ML) phylogenetic trees of *cox1* and *H3* separately using PhyML 3.1 [54]. The General time-Reversible (GTR) model and gamma distribution were selected for both markers (*cox1* and H3) based on separate test performed with jModel Test v.3 [55] and using *Zwicknia bifrons* (Capniidae) as an outgroup as described above. The trees were bootstrapped using 10,000 replications.

## Results

### Genetic diversity and DNA phylogeny

For studying the phylogeny of Nemouridae in the Japanese Archipelago, we analysed two molecular markers. *Cox1* sequences were of 658 bp length, with 247 polymorphic sites, 237 parsimony-informative sites, 10 singletons and a mean nucleotide substitution rate of 0.151. *H3* sequences were of 328 bp length, with 67 polymorphic sites, 54 parsimony-informative sites, 13 singletons and a mean nucleotide substitution rate of 0.051. No gaps were detected for either *cox1* or *H3* sequences (S1, S2 Fig). In total, for *cox1* and *H3*, we identified 128 and 68 haplotypes, respectively.

The GMYC model of *cox1* delimited 61 putative DNA-species (S1 Table). These results agreed with our 34 morphologically identified and 15 undescribed (five species of *Protonemura*, seven of *Nemoura*, one of *Indonemoura* and two of *Amphinemura*) species. Eight species (*I. nohirae*, *A. decemseta*, *A*. *zonata*, *A*. *longispina*, *A*. *megaloba*, *N. uenoi*, *N*. *chinonis* and *N*. *cf*. *cercispinosa*) showed two putative DNA-species. While *A. decemseta* showed multiple putative DNA-species (three putative DNA-species), *N*. *sanbena* and *P. kohnoae* showed two putative DNA-species in the same sampling site suggesting the presence of cryptic species. The congruence of H3 phylogenetic groups provided confirmation of DNA-based groups detected by GMYC.

We observed the genetic diversity of the species per island (Table 1). Honshu had the highest number of species (26 species), haplotype richness (63) and mean nucleotide substitution rate (average 0.027). Five species were found throughout the three Japanese islands (Honshu, Shikoku and Kyushu), i.e. *A*. *decemseta*, *A*. *zonata*, *N. cf. cercispinosa*, *N*. *chinonis* and *I*. *nohirae*, with a mean nucleotide substitution rate ranging from 0.011 to 0.126 and a total of 23 haplotypes. *N. sanbena* haplotypes were observed in two different branches in the phylogenetic tree, both within *N. cf. cercispinosa* and as an isolated branch.

**Table 1.** Regional distribution of sample size (n), haplotype richness (h) and mean nucleotide substitution rate of Nemouridae species among the three main islands in Japan, based on mitochondrial DNA (*cox1*) sequences. Total species richness was 26, 23 and 6 for Honshu, Shikoku and Kyushu, respectively.

| Genus | Species | Honshu |  |  | Shikoku |  |  | Kyushu |  |  |
| --- | --- | --- | --- | --- | --- | --- | --- | --- | --- | --- |
|  |  | n | h | Nucleotide substitution | n | h | Nucleotide substitution | n | h | Nucleotide substitution |
| *Amphinemura* | *A. bulla* | 5 | 4 | 0.006 |  |  |  |  |  |  |
|  | *A. decemseta* * | 19 | 13 | 0.014 | 9 | 5 | 0.02 | 3 | 1 | 0 |
|  | *A. dentifera* | 2 | 1 | 0 | 2 | 2 | 0.01 |  |  |  |
|  | *A. flavostigma* | 3 | 3 | 0.005 | 3 | 2 | 0.01 |  |  |  |
|  | *A. longispina* | 2 | 1 | 0 |  |  |  |  |  |  |
|  | *A. megaloba* | 4 | 2 | 0.091 | 3 | 1 | 0 |  |  |  |
|  | *A. zonata* * | 2 | 2 | 0.053 | 1 | 1 |  | 1 | 1 |  |
|  | *A. sp. n.* | 1 | 1 |  | 3 | 3 | 0.03 |  |  |  |
| *Indonemoura* | *I. nohirae* * | 9 | 4 | 0.059 | 4 | 2 | 0.01 | 2 | 1 | 0 |
| *Nemoura* | *N. akagii* | 2 | 1 | 0 |  |  |  |  |  |  |
|  | *N. cf. cercispinosa* * | 2 | 2 | 0.011 | 13 | 7 | 0.01 | 2 | 1 | 0 |
|  | *N. chinonis* * | 2 | 2 | 0.126 | 5 | 3 | 0.09 | 1 | 1 |  |
|  | *N. fulva* | 3 | 3 | 0.043 |  |  |  |  |  |  |
|  | *N. cf. hikosan* |  |  |  | 2 | 2 | 0.1 |  |  |  |
|  | *N. longicercia* | 2 | 1 | 0 | 7 | 5 | 0 |  |  |  |
|  | *N. naraiensis* |  |  |  | 2 | 1 | 0 |  |  |  |
|  | *N. ovocercia* | 1 | 1 |  |  |  |  |  |  |  |
|  | *N. redimiculum* | 1 | 1 |  | 3 | 3 | 0.01 |  |  |  |
|  | *N. sanbena* |  |  |  | 2 | 2 | 0.5 |  |  |  |
|  | *N. shikokuensis* |  |  |  | 4 | 1 | 0 |  |  |  |
|  | *N. stratum* | 2 | 2 | 0.027 |  |  |  |  |  |  |
|  | *N. speciosa* | 2 | 2 | 0.003 |  |  |  |  |  |  |
|  | *N. transversospinosa* |  |  |  | 6 | 4 | 0.01 |  |  |  |
|  | *N. uenoi* | 1 | 1 |  | 2 | 2 | 0.01 |  |  |  |
|  | *N. yakushimana* |  |  |  |  |  |  | 2 | 1 | 0 |
|  | *N. sp. n. 1* |  |  |  | 3 | 1 | 0 |  |  |  |
|  | *N. sp. n. 2* | 2 | 2 | 0.003 |  |  |  |  |  |  |
|  | *N. sp. n. 3* | 1 | 1 |  |  |  |  |  |  |  |
|  | *N. sp. n. 4* | 1 | 1 |  |  |  |  |  |  |  |
| *Protonemura* | *P. kohnoae* | 6 | 3 | 0.043 |  |  |  |  |  |  |
|  | *P. orbiculata* | 6 | 6 | 0.029 |  |  |  |  |  |  |
|  | *P. sp. n.* |  |  |  | 2 | 2 | 0.01 |  |  |  |
|  | *P. sp. n. 1* | 1 | 1 |  |  |  |  |  |  |  |
|  | *P. sp. n. 2* | 2 | 1 | 0 |  |  |  |  |  |  |
|  | *P. sp. n. 3* | 1 | 1 |  |  |  |  |  |  |  |
|  | *P. sp. n. 4* |  |  |  |  |  |  | 1 | 1 |  |

(\*) Species found on the three Japanese islands.

### Divergence dates

The Bayesian phylogenetic trees for *cox1* and *H3* showed tree topology similarity (ILD test, P = 0.8). Three clades corresponded to the three families—*Protonemura*, *Amphinemura* and *Nemoura*—whereas *Indonemoura* was divided into two clades—the Mainland China clade, clustered with *Protonemura*, and the Japanese clade (Fig 2).

The evolutionary divergence between the Nemouridae and Capniidae families was settled at 180 Ma, with a 95% HPD interval of 160 to 198 Ma, in the Jurassic geological period (Fig 2, S3 and S4 Fig). Genus-level diversifications within Nemouridae occurred in the early and middle Cretaceous. *Indonemoura* from Japan at 119.0 Ma (95% HPD, 125.8 to 100.2 Ma), *Indonemoura* from Mainland China at 112.0 Ma (95% HPD, 90.2 to 115.0 Ma), *Protonemura* at 112.7 Ma (95% HPD, 98.0 to 121.3 Ma), *Nemoura* at 107.0 Ma (95% HPD, 98.8 to 110.1 Ma) and *Amphinemura* at 80.0 Ma (95% HPD, 75.1 to 92.0 Ma). The speciation process occurred between 25 Ma (early Paleogene) and 90 Ma (late Crustaceous). Out of 35 events of speciation (i.e. nodes), 16 (45%) occurred during late Crustaceous and 19 (54%) occurred during early Paleogene, broadly overlapping with the formation time of the Japanese Archipelago (15 to 30 Ma). We observed intra-species diversification in *I. nohirae*, *A. decemseta*, *A*. *zonata*, *A*. *longispina*, *A*. *megaloba*, *N*. *chinonis*, *N. uenoi* and *N*. *cf*. *cercispinosa* (GMYC > 1 species, S1 Table). These species were divided into two clades (S5 Fig), spatially segregated into eastern and western Japan of the Fossa Magna region during the late Neogene period (20 to 22 Ma). Recent diversifications for *Nemoura* and *Amphinemura* species within either eastern or western Japanese branches were additionally revealed by TMRCA analysis of *cox1* (see Methods)*. A*. *decemseta* ranging from 3 to 3.5 Ma (95% HPD, 2.8 to 4.1 Ma); *A*. *zonata*, ranging from 3 to 4 Ma (95% HPD, 3.5 to 5 Ma); *A*. *longispina*, ranging from 3.6 to 4.5 Ma (95% HPD, 3.9 to 5 Ma); *A*. *megaloba*, ranging from 3.5 to 4 Ma (95% HPD, 2.8 to 4 Ma); *N. uenoi*, ranging from 3 to 4 Ma (95% HPD, 3.5 to 4.2 Ma) and *N*. *cf*. *cercispinosa*, ranging from 3.5 to 4.1 Ma (95% HPD, 3 to 5 Ma), for 1.5% Ma and 3.54% Ma analysis respectively.

### Phylogeographic pattern between Nemoura from Japan and North America

DNA sequences in the Japanese clade of *Indonemoura* (single species, *I. nohirae)* showed a high homology with those in the Alaskan species of *Z. columbiana* (*COI*: KM874174; >93% sequence similarity) and *Z. cinctipes* (H3: EF622600; >98% sequence similarity) based on blastn results. The ML phylogenetic trees for both *cox1* and *H3* (Fig 3) showed that the *Indonemoura* Japanese clade clustered with three North American species (*Z. columbiana*, *Z. cinctipes* and *P. delicatula*) and the *Indonemoura* Mainland China clade clustered with the East Asian Nemouridae genera (*Nemoura*, *Protonemura* and *Amphinemura*). The pairwise nucleotide substitution rate based on *cox1* between the *Indonemoura* Japanese clade and *Zapada* spp. or *P. delicatula* from North America ranged from 0.13 to 0.15, whereas a higher pairwise nucleotide substitution rate based on *cox1* of 0.26 was observed between the *Indonemoura* Japanese and Mainland China clades (Table 2).

**Fig 3.**
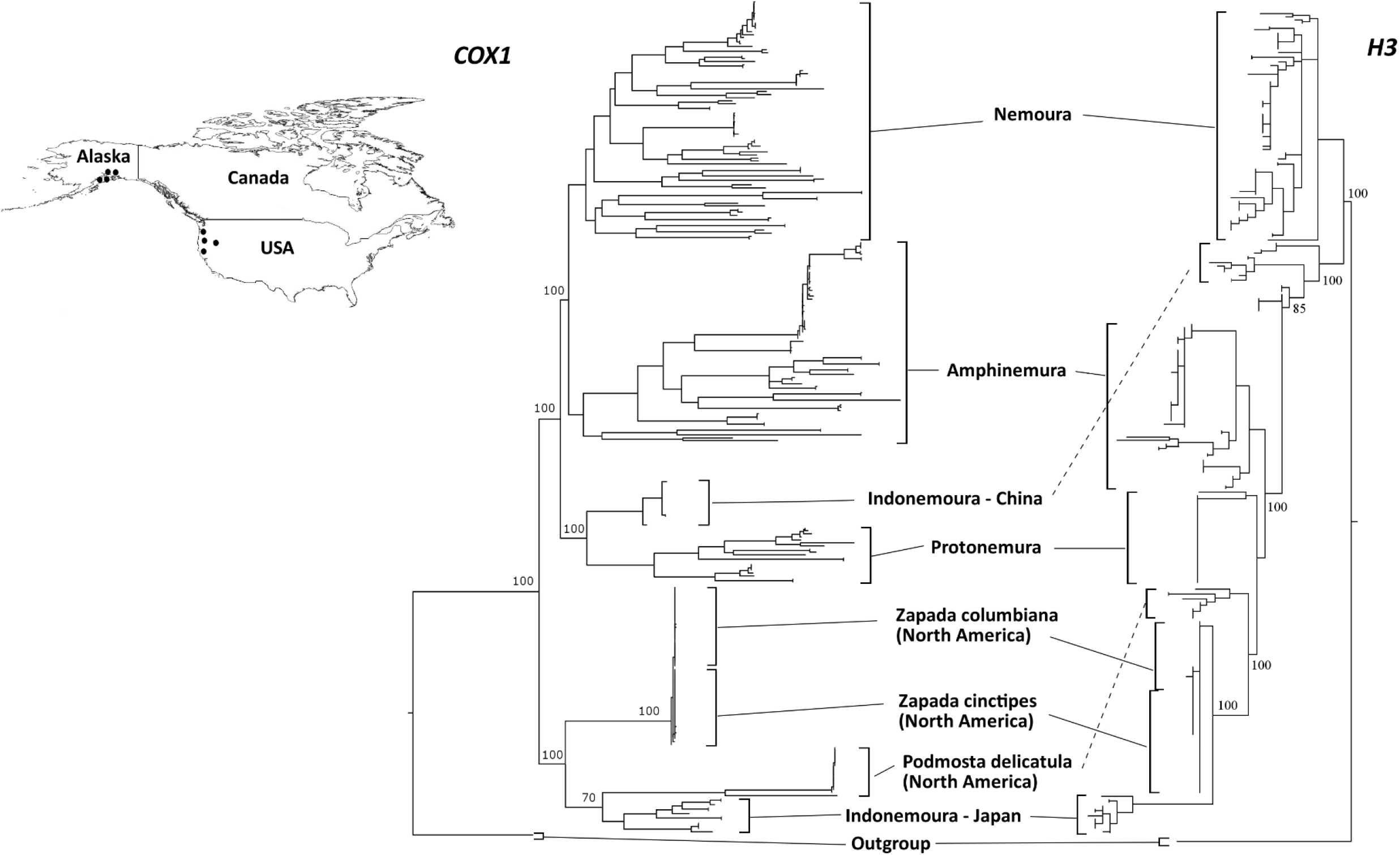
Maximum likelihood trees based on both *cox1* and *H3* markers for comparison between the East Asia Nemouridae family and three North American Nemourinae species: *Zapada cinctipes*, *Z*. *columbiana* and *Podmosta delicatula*. Inserted upper map shows sampling site locations in North America (western USA and Alaska) as black dots.

**Table 2.** Pairwise nucleotide substitution rate based on *cox1* between the East Asian Nemouridae and North American (western USA and Alaskan) species.

|  |  |  |  |  |  |  |  |  |  |
| --- | --- | --- | --- | --- | --- | --- | --- | --- | --- |
| East Asia | *Amphinemura* spp. | 0 |  |  |  |  |  |  |  |
|  | *Nemoura* spp. | 0.187 | 0 |  |  |  |  |  |  |
|  | *Protonemura* spp. | 0.197 | 0.19 | 0 |  |  |  |  |  |
|  | *Indonemoura* spp.<br>(China) | 0.213 | 0.193 | 0.197 | 0 |  |  |  |  |
|  | *Indonemoura* spp.<br>(Japan) | 0.197 | 0.183 | 0.175 | 0.260 | 0 |  |  |  |
| North America | *Zapada columbiana* | 0.178 | 0.165 | 0.182 | 0.190 | 0.145 | 0 |  |  |
|  | *Zapada cinctipes* | 0.170 | 0.154 | 0.156 | 0.179 | 0.149 | 0.133 | 0 |  |
|  | *Podmosta delicatula* | 0.202 | 0.196 | 0.191 | 0.205 | 0.135 | 0.185 | 0.201 | 0 |

## Discussion

We studied mitochondrial *cox1* and nuclear *H3* gene sequences to determine the patterns of diversification and phylogenetic relationships of species belonging to four genera of stoneflies of the Nemouridae family in the Japanese Archipelago. We estimated the divergence among *Nemoura*, *Amphinemura*, *Indonemoura* and *Protonemura* to have occurred in the early and mid-Cretaceous (around 100 Ma), which is compatible with previous studies based on fossil records [56], [57]. Our results suggested that these four genera might have dispersed and colonised different areas of the Eurasian continent—including the Japanese landmasses—when they were still connected to the Eurasian continent. Among the four genera, the diversification of *Indonemoura* occurred earlier (120 Ma) than that of the other genera (<100 Ma), suggesting that it is an ancient genus. The geological isolation of colonised areas [58], the long evolutionary time [59] and poor dispersal ability of *Indonemoura* [23], [60] might have accounted for their ancient diversification.

Based on the phylogenetic relationships of both molecular markers (*cox1* and *H3*), we observed that the three genera *Nemoura*, *Amphinemura* and *Protonemura* were monophyletic and clustered as three independent groups, as previously observed by morphological systematics [16]. However, *Indonemoura* was paraphyletic. This genus was divided into two clades corresponding to the Mainland China and the Japanese clades. Surprisingly, the Japanese clade of *Indonemoura* (single species, *I. nohirae*) clustered together with North American species (*Z. columbiana*, *Z. cinctipes* and *P. delicatula*), with a low pairwise nucleotide substitution rate (<0.15). The distribution range of these two North America genera covers North America and Eastern Asia. Previous studies suggested that their distribution could be related to the land connection (i.e. the islands) between Alaska and Eastern Asia [15]. Dispersal by island connectivity between Alaska, the Aleutian Islands, the Kamchatka peninsula and the Kuril islands has been observed in other stonefly families (for instance, *Arcynopteryx dichroa*, *Capnia nearctica*, *Mesocapnia variabilis* and *Nemoura arctica*) [61]. However, the distribution of *Indonemoura* on these islands is unknown.

The complex history of the geological formation of the Japanese Archipelago may provide a possible alternative explanation. The ancestral Japanese landmasses were located on the borders of four major tectonic plates, of which two are continental plates—the Eurasian plate and the North American plate [4] (S6 Fig). The eastern Japanese landmass was located on the North American plate, whereas the western Japanese landmass was located on the Eurasian plate [5]. The dispersal and colonisation of *Indonemoura* might have occurred from the North American plate to the Eurasian continent or vice versa (from the Eurasian continent to the North American plate) before their geographic separation in an ancient time (around 70 to 80 Ma) [62]. Dispersal events between Eurasian and Japanese landmasses are commonly reported for aquatic insects [10], [63]. Particularly, a dispersal event between North America and the Japanese Archipelago was detected by the phylogenetic relationship of the monophyletic group of caddisflies, *Palaeagapetus* spp. [9]. However, no prior studies have observed speciation events of aquatic insects associated with geological events that occurred in ancient times (>12 Ma). Our result suggests an ancient divergence time and a distribution pattern of *Indonemoura*, consistent with a hypothesis of an ancient colonisation influenced by the connection of the Japanese landmass with the North American plate in the Eurasian continent.

Nemouridae species diversification, as has been observed in other species of aquatic insects, such as beetles [10], caddisflies [9], water bugs [14] and mayflies [8], [13], was also observed to be affected by the geological formation of the Japanese Archipelago. The diversification of the Nemouridae species occurred during the Paleogene period (<50 Ma). This geological period is consistent with the movement of landmasses (S6 Fig) about 70 Ma ago [4] and the active geological formation of the Japanese Archipelago around 20 Ma ago [5], which could be the cause of the Nemouridae diversification, as previously reported for the mayfly *Dipteromimus flavipterus* (35 Ma) [2].

*Indonemoura nohirae* is the single species of *Indonemoura* on the Japanese Archipelago [25], [26]. The morphology of their terminalia resembles that of *Protonemura* rather than of *Indonemoura*, but the characteristic gill formula justifies their taxonomical classification in *Indonemoura* [25], [26]. To date, there are 24 *Indonemoura* species from China [16], [24] and 30 species belonging to the Himalayan and Oriental regions in East Asia [15], [20]. These species are morphologically different from *I*. *nohirae* in Japan [15], [16], [20], [24], [25], [26]. We hypothesise that the *Indonemoura* species of East Asia could be forming separate phylogenetic clades clustered by geographical regions. For the hypothesis testing, further collection of molecular data on *Indonemoura* from wider areas such as Northeast China, Southeast China, Mongolia, Russia and other countries in Asia is needed in future studies.

Eight species (*I. nohirae*, *A. decemseta*, *A*. *zonata*, *A*. *longispina*, *A*. *megaloba*, *N*. *chinonis*, *N. uenoi* and *N*. *cercispinosa*) showed interesting patterns of intra-species separation into two genetic groups corresponding to eastern and western areas of the Fossa Magna region of Honshu Island (S5 Fig). Honshu is the centre of insect biodiversity [10]; apart from its extensive territorial space, it is the main island with a geological history [3], [4], [5]. We found supporting evidence on the genetic diversity of these eight species. We found a larger mean nucleotide substitution rate and haplotype number in the Honshu region than in other islands (Table 1). The mean nucleotide substitution rate and haplotype diversity are indications of biodiversity [64], which could lead to evidence of speciation [65]. Out of eight species, the diversification of six species (*A. decemseta*, *A*. *zonata*, *A*. *longispina*, *A*. *megaloba*, *N. uenoi* and *N*. *cercispinosa*) occurred during the late Neogene period (20 to 22 Ma). This event corresponded with the double-door (i.e. the union of eastern and western Japan; S6 Fig) geological model and the formation of the Itoigawa-Shizuoka tectonic line (ISLT) at around 20 Ma [5], [66]. The speciation of aquatic insects was often observed to be influenced by these two geological events [2]. Additionally, species diversification—from eastern or western Japan of the Fossa Magna region—showed recent diversification events (3 to 5 Ma) corresponding with the formation of the small islands in northeastern or southwestern edge areas of Japan (Fig 1). The northeastern islands created land bridges between the Japanese Archipelago and China or Korea, whereas the southwestern islands connected Taiwan or the Philippines with the Japanese Archipelago [5], [66]. This connectivity promoted immigration events in Japan that might have contributed to the formation of the current genetic diversity, as previously observed in mayflies [13] and beetles [10].

The evolutionary divergence of the Nemouridae family was promoted by the complex geological formation of the Japanese Archipelago. Despite the different evolutionary rate of both molecular markers, Bayesian analysis found congruence between both markers; however, failed to find congruence with their morphological taxonomy. The main morphological character used for identification of adult stoneflies species is its genital morphology. The evolution of genital morphology is, however, governed by within-population sexual selection rather than environmental or geological history of the locations [67]. Conversely, the genetic variation of natural populations has been observed to be directly associated with environmental [68] and geological variations [2]. Therefore, the genetic variation could reflect an independent course in the evolutionary history of *Indonemoura* than do the morphological characters used for their taxonomy. However, we detected that *N*. *sanbena* shared haplotypes from different lineages, revealing a possible introgression or incomplete sorting of ancestral polymorphisms [10]. This is an often reported phenomenon in stoneflies [40], [69], [70], which remains as unresolved species. Resolving the problems between the process of evolution of morphological characters and the genetic variation within species will improve our future understanding of the origin of the species and the local species distribution.

Finally, our inference of divergence time was based on the coalescent simulation approach. Despite the frequent use of this approach, a biased sampling of lineages and extreme state-dependent molecular substitutions rate heterogeneity are known to potentially cause erroneous inference of divergence time [71]. Therefore, combining node calibrations generated by more than one calibration analyses is recommended [71], [72]. A cautious method such as the combined uses of fossil records and biogeographic ages as employed in our analysis may minimise the risk of such erroneous inference.

## Acknowledgments

We thank the following colleges for contributing Nemouridae specimens for our analysis: Akatsuki Yoshinari, Kobayashi Kyota, Weihai Li and Boris C. Kondratieff. We also acknowledge anonymous reviewers for constructive comments on an earlier version of the manuscript.

## Supplementary information

**S1 Table.** Location information of samples of East Asia Nemouridae. Numbers of individuals (N), presences of male (M), female (F) and imago (im), DNA-species delimitation (GMYC).

**S1 Fig.** Multiple alignment of *cox1* region for the 54 Nemouridae genotypes.

**S2 Fig.** Multiple alignment of *H3* regions for the 54 Nemouridae genotypes.

**S3 Fig.** *Cox1* Bayesian trees using three types of node calibration. A – fossil, B – island formation and D – TMRCA.

**S4 Fig.** *H3* Bayesian tress using two types of node calibration. A – fossil and B – islands formation.

**S5 Fig.** Concatenated Bayesian phylogeny (*cox1* + *H3*) for East Asian Nemouridae family enlarging intra-species diversification in *I. nohirae*, *A. decemseta*, *A*. *zonata*, *A*. *longispina*, *A*. *megaloba*, *N*. *chinonis*, *N. uenoi* and *N*. *cf*. *cercispinosa* (GMYC = 2 species).

**S6 Fig.** Putative formation of the Japanese Archipelago [2], [3], [4], [5]. (A) Around 30 to 130 Ma, the Japanese landmasses were located in two major tectonic plates from the Eurasian continent. (B) Around 15 to 30 Ma, the Japanese landmasses began to separate from Eurasia and the North American Plates began to separate from the Eurasian continent, and remained separated by a sea zone called Fossa Magna—a geological event called double-door. (C) Current map of the Japanese Archipelago in East Asia, where the names of the main four Japanese islands and the two tectonic lines are shown.

